# Mind the drift - improving sensitivity to fMRI pattern information by accounting for temporal pattern drift

**DOI:** 10.1101/032391

**Authors:** Arjen Alink, Alexander Walther, Alexandra Krugliak, Jasper J.F. van den Bosch, Nikolaus Kriegeskorte

## Abstract

Analyzing functional magnetic resonance imaging (fMRI) pattern similarity is becoming increasingly popular because it allows one to relate distributed patterns of voxel activity to continuous perceptual and cognitive states of the human brain. Here we show that fMRI pattern similarity estimates are severely affected by temporal pattern drifts in fMRI data – even after voxel-wise detrending. For this particular dataset, the drift effect obscures orientation information as measured by fMRI pattern dissimilarities. We demonstrate that orientation information can be recovered using three different methods: 1. Regressing out the drift component through linear modeling; 2. Computing representational distances between conditions measured in independent imaging runs; 3. Crossvalidation of pattern distance estimates. One possible source of temporal pattern drift could be random walk like fluctuations — physiological or scanner related — occurring within single voxel timecourses. This explanation is consistent with voxel-wise detrending not alleviating pattern drift effects. In addition, this would explain why crossvalidated pattern distances are robust to temporal drift because a random walk process is expected to give rise to non-replicable drift directions. Given these findings, we recommend that future fMRI studies take pattern drift into account when analyzing pattern similarity as this can greatly enhance the sensitivity to experimental effects of interest.

## 1. Introduction

Multivariate analysis of functional magnetic resonance imaging (fMRI) data allows one to relate distributed patterns of activity to perceptual and cognitive states of the human brain. As pattern-information techniques are gaining popularity, it is important to identify stimulus-unrelated factors influencing fMRI patterns in order to reduce nuisance variation, avoid confounds, and make results interpretable. In this study we investigate the effect of stimulus unrelated temporal drifts, which has recently been shown to profoundly alter fMRI patterns evoked by a diverse set of visual images (Henriksson et al., 2015; Kay et al., 2008). In particular, we investigate the consequences of pattern drift on fMRI pattern dissimilarity analysis (Kriegeskorte et al., 2007; Kriegeskorte et al., 2008; Kriegeskorte & Kieviet, 2013).

In the present study, we show that temporal pattern drift also affects well-documented fMRI patterns evoked in V1 by visual orientation (Kamitani & Tong, 2005; Haynes & Rees, 2005). Specifically, we find that the size of orientation effects on fMRI patterns in V1 is dwarfed by the effect of temporal pattern drift. This effect occurs regardless of high-pass filtering and detrending of single voxel timecourses, which suggest that conventional univariate temporal preprocessing steps (Tanabe et al., 2002) do not remedy the observed pattern drift. We then demonstrate that the drift confound can be alleviated using three different methods: 1. Regressing out the drift component through linear modeling; 2. Computing representational distances between conditions measured in independent imaging runs; 3. Crossvalidation of pattern distance estimates.

## 2. Material and methods

We analyzed fMRI response patterns elicited by visual orientation stimuli in early visual areas. The data have previously been analyzed in Alink et al. (2013), where a more detailed description of the stimuli and design can be found.

### 2.1 Experimental design and task

#### 2.1.1 Experimental design

The experimental paradigm in Alink et al. (2013) was devised so as to classify different orientations of low-level visual stimuli. Four stimulus types were presented, each comprising two orthogonally oriented stimuli (see Appendix Fig. A1). Stimulus types were gratings, spirals, and versions of both in which the image had been divided into a log-polar checkerboard array of patches and half the patched had been swapped between the stimuli.

**Figure 1:**
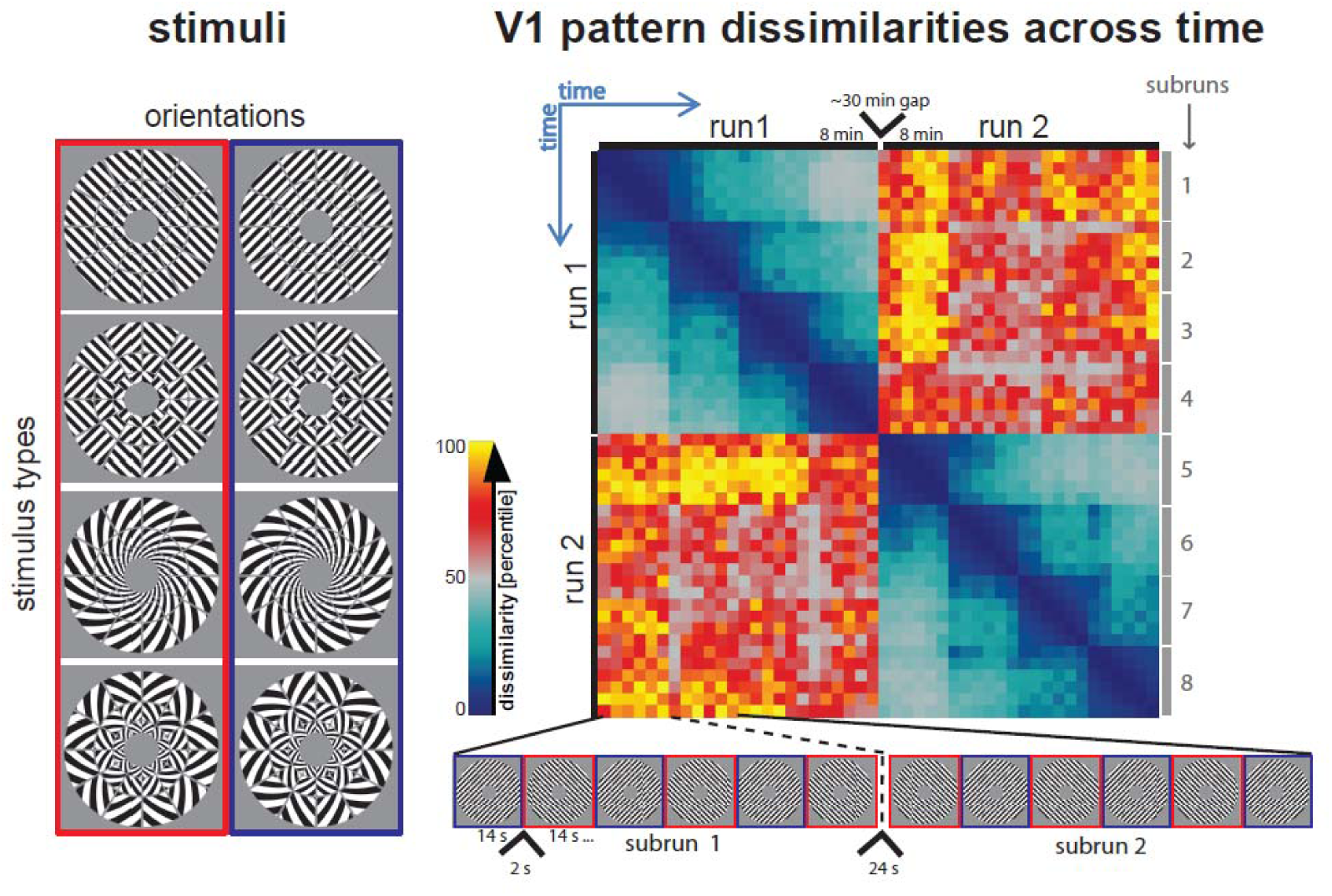
Experimental design and its relation to the chronologically ordered representational dissimilarity matrix. *Left* the four stimulus types and their orientations presented during the experiment. *Right* the chronologically ordered representational dissimilarity matrix (RDM) and its relation to the temporal structure of stimulus presentation. Mind that the RDM shown is the average RDM across all four stimulus types and subjects.

Figure 1 illustrates the temporal sequence of the stimulus presentation. Stimuli were presented in a single fMRI session with eight scanner runs, each of which lasted eight minutes. In each run, both orientations of one stimulus type were shown (e.g. gratings orientation one and orientation two). For each stimulus type, two runs were recorded. Each run consisted of four equally long subruns comprising six stimulus trials: three trials for each orientation and alternating orientations across trials, with the leading orientation alternating across subruns. Trial duration was 14 s. Each trial contained phase-randomized versions of a single orientation. During a stimulus block, 28 phase-randomized versions of the orientation were presented at a frequency of 2 Hz. The stimulus duration was 250 ms, followed by an interstimulus interval (ISI) of 250 ms. The 28 stimuli had random spatial phases, uniformly distributed between 0 and 2π. Stimulus blocks were separated by 2-s fixation periods and subruns by 24-s fixation periods. A small task-related ring around the fixation dot was visible throughout the entire run.

#### 2.1.2 Subjects and task

18 healthy participants (13 female) with normal or corrected-to-normal vision underwent scanning. During both the main experiment and retinotopic mapping a dot was presented at the center of the screen (diameter: 0.06° visual angle) which the participants were instructed to fixate continuously. The fixation dot was surrounded by a black ring (diameter: 0.20°, line width: 0.03°) with a small gap (0.03°) that randomly alternated between the left and the right side — on average once per three seconds and the minimum time between a side-switch was one second. The participants were instructed to continuously indicate whether the gap was left or right by holding down the left button with the right index finger or the right button with the right middle finger, respectively. The purpose of this task was to enforce fixation and to draw attention away from the stimuli.

### 2.2. MRI measurement and analysis

#### 2.2.1 MRI measurements

MRI images were acquired on a 3T Siemens Trio using a 32-channel head coil. During the main experiment, each functional run acquired 252 volumes containing 31 slices using an EPI sequence (TR=2000 ms, TE=30 ms, flip angle=77°, voxel size: 2.0 mm isotropic, field of view: 205 mm; interleaved acquisition, GRAPPA acceleration factor: 2). During the retinotopic mapping, we acquired 360 volumes using the same EPI sequence. Additionally, high-resolution (1 mm isotropic) T1-weighted anatomical image were obtained for each subject using a Siemens MPRAGE sequence.

#### 2.2.2 Pre-processing

Functional and anatomical MRI data were preprocessed using the Brainvoyager QX software package (Brain Innovation, v2.4). We discarded the first two EPI images for each run to prevent T1 saturation effects in the estimation of the response pattern baseline. Pre-processing comprised slicescan-time correction, 3D head-motion correction and temporal high-pass filtering removing frequencies below 2 cycles per run (frequencies lower than .004Hz). The functional images for all subjects were then aligned with the individual high-resolution anatomical image and transformed into Talairach space (Talairach & Tournoux, 1988) as a step toward cortex-based analysis in BrainVoyager. After automatic correction for spatial inhomogeneities of the anatomical image, we created an inflated cortex reconstruction for each subject. All ROIs for V1 were defined in each individual subject’s cortex reconstruction and projected back into voxel space.

#### 2.2.3 Delineation of V1 through retinotopic mapping

In order to define V1, we presented dynamic grating stimuli designed to optimally drive early visual cortex. These stimuli were based on a log-polar array, but without the grout lines and with 20 patches per ring. Each patch contained rectangular gratings with a spatial period of one third of the patch’s radial width. Grating orientation and phase was assigned randomly to each patch. Over time, the phase of the gratings increased continuously (1 cycle per second) resulting in continuous motion in each patch (in different directions). In addition, the orientation of the grating increased in steps of π/6, once each second resulting in motion direction changes within patches over time. We used five such stimuli, driving different parts of the retinotopic maps in early visual cortex: (1) a horizontal double-wedge stimulus, spanning a polar-angle range of +/-15° around the horizontal meridian, (2) a vertical double-wedge stimulus of the same kind, (3) a stimulus that covered the region driven by the main-experimental stimulus (1.50°-7.04° eccentricity), (4) a 0.5°-wide ring peripherally surrounding the main-experimental stimulus annulus (7.04°-7.54° eccentricity), and (5) a 0.5°-wide ring inside the annulus (1.00°-1.50° eccentricity). Stimuli were presented in 6-s blocks. This block length was chosen to balance temporal concentration (which increases design efficiency for long blocks due to hemodynamic buildup) and stimulus adaptation (which reduces design efficiency for long blocks due to reduced neuronal responses). The five dynamic stimuli and 6-s fixation periods were all presented 20 times each in a random sequence over a single run lasting 12 min.

An ordinary least squares general linear model (GLM) was fitted to the retinotopic mapping data, with five predictors for the five dynamic grating stimuli based on convolving boxcar functions with the hemodynamic response function as described by Boynton et al. (1996). Activation t-maps for each stimulus type were projected onto polygon-mesh reconstructions of individual subjects’ cortices. We determined the borders of V1 based on cortical t-maps for responses to vertical and horizontal double-wedge stimuli (Sereno et al., 1995). We defined ROIs for V1 as the portion of V1 that was more active when presenting the dynamic grating stimulus covering the main-experimental annulus as compared to central and peripheral stimulation (average numbers of voxels for V1: 1126, 1242 and 1031, respectively, with left and right hemispheres combined).

#### 2.2.4 Estimation of fMRI response to oriented stimuli

Pre-processed fMRI timecourses and subject-specific V1 coordinates were imported into Matlab (The Mathworks, Natick, MA, USA) using Neuroelf v0.9c (http://neuroelf.net). For each 14-s block, a response pattern was estimated with a GLM using ordinary least squares. Before univariate modeling, the timecourse data were converted to percent signal change. An individual GLM was estimated for each stimulus type, containing 48 stimulus predictors (2 runs × 4 subruns × 6 blocks). The predictor time courses were computed using a linear model of the hemodynamic response (Boynton et al., 1996). In addition to the stimulus predictors, for each run the model contained six 3D head motion predictors and one run intercept. For each voxel, we then performed a GLM fit to obtain a response-amplitude for each of the 48 blocks. Beta response estimates were then multivariately normalized by an estimate of the voxel variance-covariance matrix (Walther et al., under revision). We used a covariance estimator with optimal shrinkage (Ledoit and Wolf, 2004) toward a diagonal covariance matrix. These noise-normalized beta weights were then used for subsequent analyses.

#### 2.2.5 Classification of stimulus identity

In the original study (Alink et al., 2013), we estimated responses based on one predictor for each stimulus type and orientation per subrun — in contrast to the single block estimates used here. To test if this approach leads to similar decoding accuracies as the original study we replicated the results from our previous study on orientation effects in V1 using the same classifier, a linear support vector machine (SVM). To keep results consistent with the results in this manuscript, decoding was performed on multivariately normalized beta coefficients (whereas in the original study classification was done on *t* values). Like in our previous study, SVM was trained on seven subruns and crossvalidated on the remaining held-out subrun, resulting in eight classification folds. Classification accuracies were then averaged across folds and subjects. Results were in overall agreement with those published previously (Figure A1).

### 2.3 Representational similarity analysis of time-ordered response patterns

In the introduction, we pointed out that fMRI patterns contain contributions from temporally correlated nuisance factors. In order to assess the relationship between the temporal proximity of two given conditions and their pattern similarity, we ordered the 48 patterns of each run by the sequence in which they had been presented to the subject in the scanner. On those patterns, we computed a 48 × 48 representational dissimilarity matrix (RDM) using Pearson correlation distance. The RDMs shown in figures 1 to 4 always depict the average across stimulus types and subjects. To estimate a two-dimensional representation of the RDM, we employed non-classical multidimensional scaling (MDS) with optimization criterion metric stress (Kruskal, 1964).

### 2.4 Estimation of the orientation information index *δ*

To quantify if there was significant orientation information in the similarity structure in V1, we computed the mean of all dissimilarities between stimuli with identical orientations 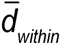 and the mean of all dissimilarities between stimuli with different orientations 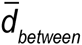 and computed an *orientation information index δ* as the difference between them:

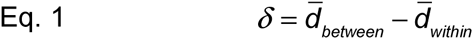

If *δ* is significantly greater than 0, this means that the dissimilarity between patterns elicited by different orientations is greater than the dissimilarity between identical orientations, indicating orientation information. A *δ* that is significantly smaller than 0 indicates that patterns evoked by identical stimuli are more similar than those evoked by stimuli with different orientations. Therefore, the finding of *δ* being significantly smaller than 0 is uninterpretable. *δ* was computed for each RDM of each subject and stimulus type. For each stimulus type, we then tested if *δ* was significantly above or below zero by a *t* test (p<0.05) across participants.

### 2.5 Recovering orientation information from fMRI pattern drift

We already alluded to the confounding influence of drifts between temporally adjacent pattern estimates. Here, we introduce three methods to control for pattern drift distortions in the similarity structure: 1. Regressing out the drift component through linear modeling; 2. Computing representational distances between conditions measured in independent imaging runs; 3. Computing crossvalidated distance estimates (Nili et al., 2014; Walther et al., under revision).

#### 2.5.1 Linear modeling of pattern drift

A simple way of allaying temporal distortions in the similarity structure is to estimate their contribution to the overall dissimilarity variance and to take out this variance component. We can determine the weight of this contribution by applying a general linear model to the dissimilarity matrix by which we model the temporal drift. By default, this model contains an intercept with weight *β*_0_ (meaning the regressor has the same value for all dissimilarities and therefore models the overall dissimilarity score) and a drift regressor **dr**_k_ with weight *β_k_*. The drift regressor predicts any given dissimilarity value in the measured RDM as a function of the time elapsed between its two associated conditions in the fMRI experiment. Since drift distortions are predominantly time-dependent, this regressor will by proxy measure the drift dissimilarity variance component visited onto the RDM.

To determine the best fitting drift function describing the measured RDM, we defined 24 polynomial drift models with increasing degrees n, where the 1^st^ degree polynomial only contains a linear drift predictor while the 24^th^ degree model has 24 drift-related weights:

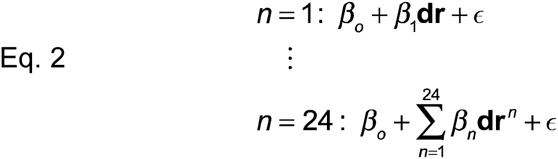
where ∊ are the model residuals. To model the contribution of pattern drift to the dissimilarity structure, we estimated the fit for each of the 24 models to the RDM. Model fits were performed using ordinary least squares.

To determine the best-fitting drift model, we computed a *drift-velocity estimate*. We defined the drift-velocity estimate as the average difference between dissimilarity residuals of the same kind (same or different orientation) of each subrun. Hence, the index measures the consistency of the dissimilarity over time in the subruns. If the fMRI patterns are drift-stable (i.e. consistently reinstated in independent blocks), the index will be close to zero. If the fMRI patterns are drift-perturbed, the index will be either larger or smaller than zero, depending on the direction of the effect. For each subject, we computed the drift velocity estimate for each drift model. *δ* was then computed on ∊ of the lowest-degree model with a drift velocity estimate that was not significantly different from zero.

#### 2.5.2 Computing the between-run correlation distance

Another method to recover dissimilarity values from pattern drift is to compute the distance measure between fMRI patterns from two independent repetitions of the same stimulus set. In fMRI, such independent data are provided by functional imaging runs in between which scanning is stopped. For two given conditions a and b, the Pearson correlation can be computed as the cosine of the angle between the mean-centered estimated activity pattern of condition a of run one, 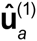, and condition b of run two, 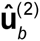

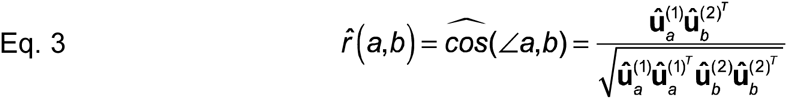

And 1-r is the correlation distance between a and b.

The estimated fMRI patterns can be assumed to be composed of two additive pattern components: a true condition-specific pattern, e.g. **u***_a_*, and a run-specific noise pattern, e.g. ∊*_a_*, which includes stimulus-unrelated pattern drift. For two conditions belonging to their respective run one and two, the fMRI pattern estimates of a and b obtain as

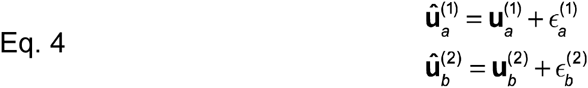

While the truly condition-related pattern component will be stably reinstated in repeated measurements, noise patterns can be assumed independent between different functional runs, since these runs are independent measurements themselves, hence are random fluctuations between them. Note that this does not rule out the possibility that the generating noise processes may be very similar in individual runs, which may give rise to temporally correlated fMRI noise within each imaging run, accounting for noise drifts between temporally adjacent conditions.

Substituting the estimated activity patterns in Eq. 3 for their components in Eq. 4 obtains as

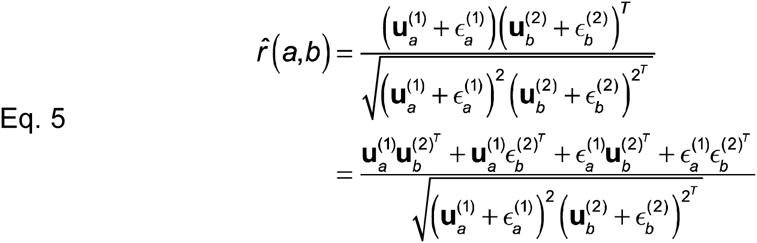

As alluded to (Eq. 4), each noise pattern is independent to any other activity pattern belonging to a different run. Therefore, the expected value of 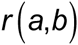 is

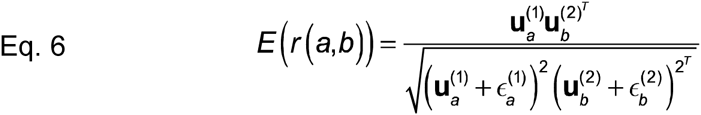

Therefore, the expected value of the correlation between a and b will reflect the true covariance between a and b, if a and b come from independent runs. Note that in the denominator, the variance of a and b are still noise-biased because they are confined to their respective runs, hence the error terms are retained.

For each condition pair, this procedure yielded two between-run correlation distance estimates (note that 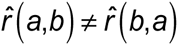 because the variances are different in run 1 and 2), which were subsequently averaged. For each subject and stimulus type, all pairwise between-run dissimilarities were then assembled in a RDM on which the orientation information index *δ* was computed.

#### 2.5.3 Crossvalidated Mahalanobis distance estimate

A third way of computing a drift-corrected dissimilarity measure is by crossvalidating the distance in independent data. Like between-run dissimilarities (see section 2.5.2), crossvalidated distance estimates (Walther et al, under revision; Nili et al, 2014) are not affected by artificially blown-up pattern covariances. In addition to that, they are bound to an interpretable zero point, meaning they are ratio-scale. Moreover, while between-run dissimilarities only restore the noise-unbiased between-condition covariance (see Eq. 5), crossvalidated distance estimates also preserve the true pattern variances in the expected value (see 7.1 in the appendix)

Unlike the conventional Pearson correlation coefficient, a crossvalidated correlation estimate is not bounded between -1 and 1 anymore: as the voxel patterns of condition a and b belong to different runs, they may vary substantially in voxel variance. Therefore, although the resulting crossvalidated correlation estimate will come from a distribution around the true correlation value, the estimate need not conform to the boundaries of the Cauchy-Schwarz inequality and can exceed the range of the Pearson correlation. This makes the value harder to interpret and does not comply with the definition of the correlation distance, which is one minus r.

Instead, we computed the crossvalidated squared Mahalanobis distance estimate between all possible condition pairs (Walther et al., under revision):

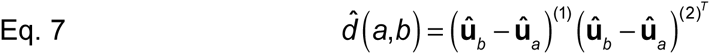

Before calculating the dissimilarity measure, we applied voxel mean subtraction and voxel variance normalization to the fMRI patterns. This is sensible because both normalizations are also implicitly carried out by the correlation distance, and make the squared Euclidean distance proportional to the correlation distance (Nili et al., 2014).

We computed crossvalidated squared Mahalanobis distance estimate RDMs of all pairs of conditions for each subject and stimulus type, from which *δ* was then obtained.

## 3. Results

### 3.1 Temporal drift severely distorts fMRI pattern geometry

Two visual features of the temporally ordered RDM (see figure 2a) clearly stand out: a prominent dark blue band centered about the diagonal and the yellow-red colored squares for dissimilarities across runs. The dark blue band along the diagonal indicates that fMRI patterns in close temporal proximity are more similar to each other than any other fMRI patterns. In order to quantitatively determine the prominence of this effect we computed Kendall’s *τ_a_* (Nili et al., 2014) between fMRI pattern correlation distances and temporal proximity of the stimuli in the experimental sequence — constrained to within run pattern dissimilarities (figure 2c). We observed an average *τ_a_* of 0.41 (t_17_=30.08, p<0.001), indicating a prominent linear temporal drift component to the dissimilarity structure. The fact that *τ_a_* between orientation differences (1 for different orientation and 0 for same orientation) and pattern dissimilarities was -0.005 (t17=-12.55, p<0.001, section 3.3 explains why this correlation is negative) highlights that temporal drift has a much greater impact on fMRI pattern similarity than the experimental effects of interest. The prominence of the drift is also illustrated by the two-dimensional representation of the RDM obtained by multidimensional scaling (MDS) (figure 2b), where temporally adjacent patterns are linked by a gray line, revealing that response patterns measured in temporal proximity are much more similar than response patterns elicited by the same stimulus.

**Figure 2:**
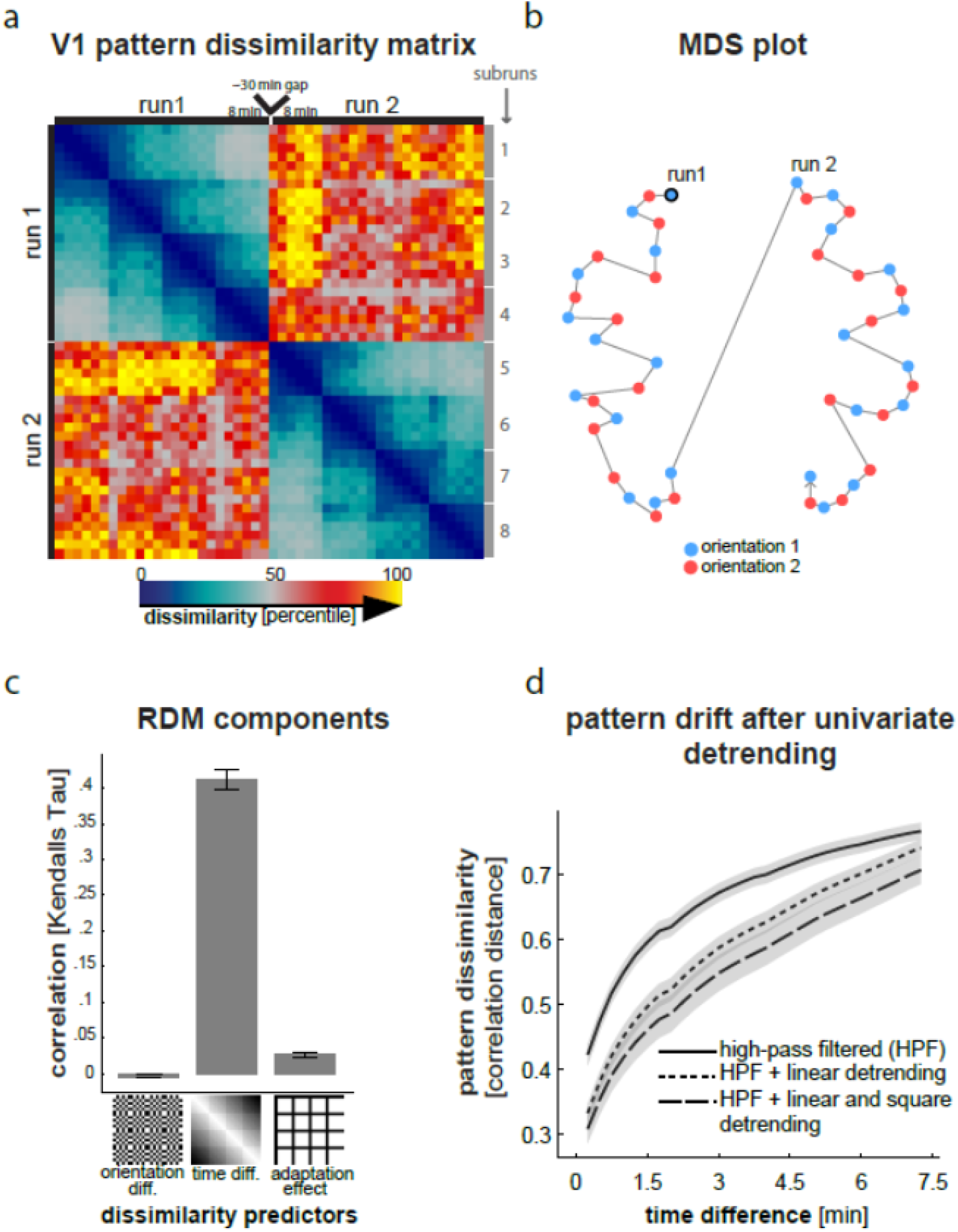
Impact of temporal drift on fMRI pattern similarity. *a*. same chronologically ordered RDM as shown in figure 1 b. 2D multidimensional scaling plot illustrating the relative impact of orientation, temporal proximity and run effects on pattern similarity c. results of a correlation analysis investigating the extent to which pattern dissimilarity is affected by orientation, temporal drift and adaptation d. line plot indicating that pattern dissimilarity increases as a function of inter-pattern time difference. The solid line depicts the drift after temporally high-pass filtering at .004Hz – which corresponds to the data used in all other analyses. The other lines depict how additional univariate linear and square detrending affect temporal pattern drift.

The second feature — the yellow-red colored squares for dissimilarities across runs — indicates that fMRI patterns tend to be very different when measured during different runs. The average between-run correlation distance was 0.96, whereas the average within-run correlation distance was 0.71. Because runs are on average 29.77 minutes apart (standard deviation 13.86 minutes), it is likely that this effect relates to a temporal drift effect. An additional factor, however, is the fact that the scanner is switched off and on between runs which might also contribute to greater between than within run pattern dissimilarities. In addition, confound means were estimated separately for each run which should further increase pattern correlation distances between runs.

In the above mentioned RDM correlation analysis, we modeled the temporal drift as proportional to the temporal separation of the two blocks whose response patterns are being compared. Plotting fMRI pattern correlation distances as a function of temporal proximity, however, indicates that the drift effect on pattern dissimilarity saturates with larger temporal separations (figure 2d, solid line). The dissimilarity between fMRI patterns is most strongly affected by temporal drift when fMRI patterns are in close temporal proximity. This temporal profile was found to be highly consistent across participants (see shaded standard error ranges in figure 2d around the solid line).

### 3.2 Temporal pattern drift is not remedied by voxelwise detrending

One possible source of pattern drift could be low frequency changes in single voxel time-courses. This, however, is unlikely given that we have high-passed filtered our data at .004Hz. In addition to high-pass filtering we have also included voxelwise linear and combined linear and square detrending as preprocessing steps. Both types of detrending led to an overall reduction of pattern dissimilarities (figure 2d). The slope of the drift, however, was found to be unaffected by voxelwise detrending.

### 3.3 Impact of repetition suppression on fMRI pattern geometry

Another more subtle visual feature of the chronologically ordered RDM are the faint dark blue lines overlapping with rows and columns corresponding to the first stimulus presentations within subruns (figure 2a). These lines indicate that fMRI patterns evoked by first stimuli in subruns are more similar to all other fMRI patterns than those evoked by later stimulus presentations within a subrun. The *τ_a_* correlation between the RDM and the corresponding predictor RDM was 0.03 (t17=7.28, p<.001, figure 2b). We attribute this effect to first stimuli within each subrun being least affected by repetition suppression (Grill-Spector et al., 2006). This gives rise to larger responses for the first stimulus in a subrun which should increase these patterns’ signal to noise ratios (SNR) relative to the other patterns. This would explain the enhanced pattern similarities between these patterns and all other patterns.

### 3.4 Drift-uncorrected analysis of orientation pattern similarity leads to uninterpretable results

One troubling finding — touched upon in 3.1 — is that there is a significant negative correlation between pattern dissimilarity and orientation difference (*τ_a_*=0.41, t_17_=30.08, p<0.001, figure 2c). This indicates that patterns evoked by a stimulus with a different orientation are consistently more similar to each other than patterns evoked by the same stimulus. We know, however, that the patterns do carry information with regard to orientation based on the results of our previous study (see Fig. A1 in the Appendix and Alink et al., 2013). The main reason for this discrepancy is the fact that temporal proximity and stimulus orientation were confounded in the experimental design: due to the alternating fashion of presentation, temporally adjacent blocks always had opposite orientations within each subrun. Because temporal proximity strongly reduces pattern dissimilarity (figure 2d), this confound leads to the observed lower average distance for patterns with different orientations.

This shows that simply comparing within- and between-orientation pattern dissimilarities can lead to uninterpretable results if stimulus sequence is not randomized. In the next two sections, we will describe analysis methods that remove the effects of temporal drift and recover fMRI pattern information about stimulus orientation.

### 3.5 Orientation information can be recovered by regressing out drift effects from pattern dissimilarities

We have seen that temporal pattern drift can render a naive comparison of average within- and between-condition pattern dissimilarities unintepretable. Here we describe how one can alleviate this problem by regressing out temporal-drift-related pattern variance. To this end, we performed polynomial regression using 24 different drift models with 1 to 24 degrees (figure 3a, see section 2.5.1 for details). For each model, we obtained the RDM residuals e. In order to test whether e was unaffected by the effect of temporal drift we computed the drift velocity estimate of each model (figure 3b, see section 2.5.1 for details). A polynomial drift model with five degrees was found to be the most parsimonious model that removed temporal drift (figure 3b).

When computing orientation information (*δ*) based on this model’s residual RDM we found that *δ* was significantly greater than zero (average *δ* across subjects and stimuli was 0.0074; p<0.0001), suggesting the presence of pattern orientation information. In addition, *δ* across stimulus types (figure 3c) was found to be qualitatively similar to that obtained in our previous study using SVM classification (Alink et al., 2013 and Fig A1). Therefore, it appears that temporal drift effects can be regressed out at the RDM level and that this increases sensitivity to fMRI pattern effects of interest.

**Figure 3:**
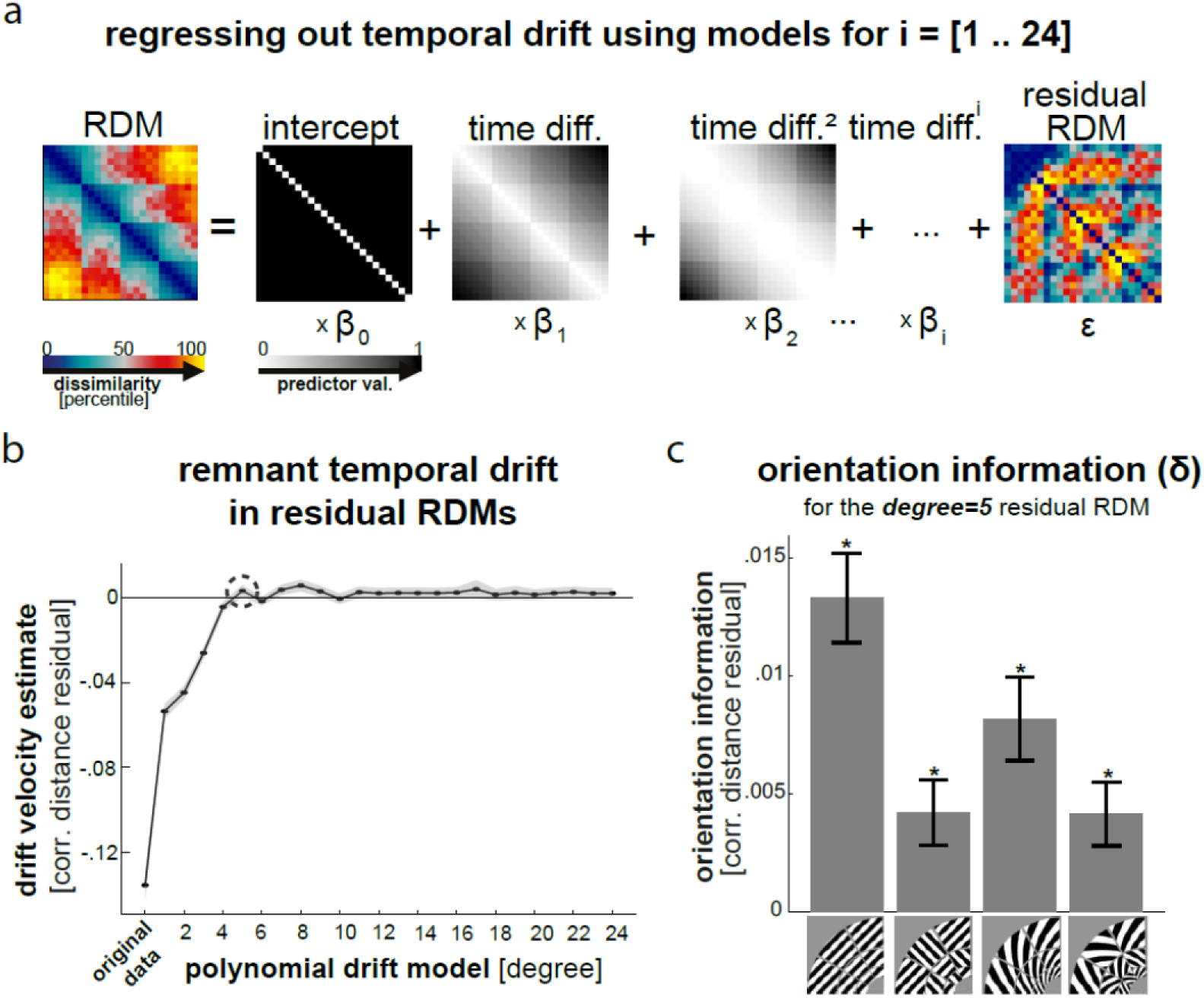
Removing drift effects by linear modeling of pattern dissimilarities as a function of stimulus time difference. *a*. exemplary illustration of the polynomial models used with degrees ranging from 1 to 24 *b*. line plot showing the drift velocity index - the average difference between dissimilarity residuals of the same kind (same or different orientation) within each subrun - as a function of model degree *c*. bar graph depicting recovered orientation information for each stimulus type.

**Figure 4:**
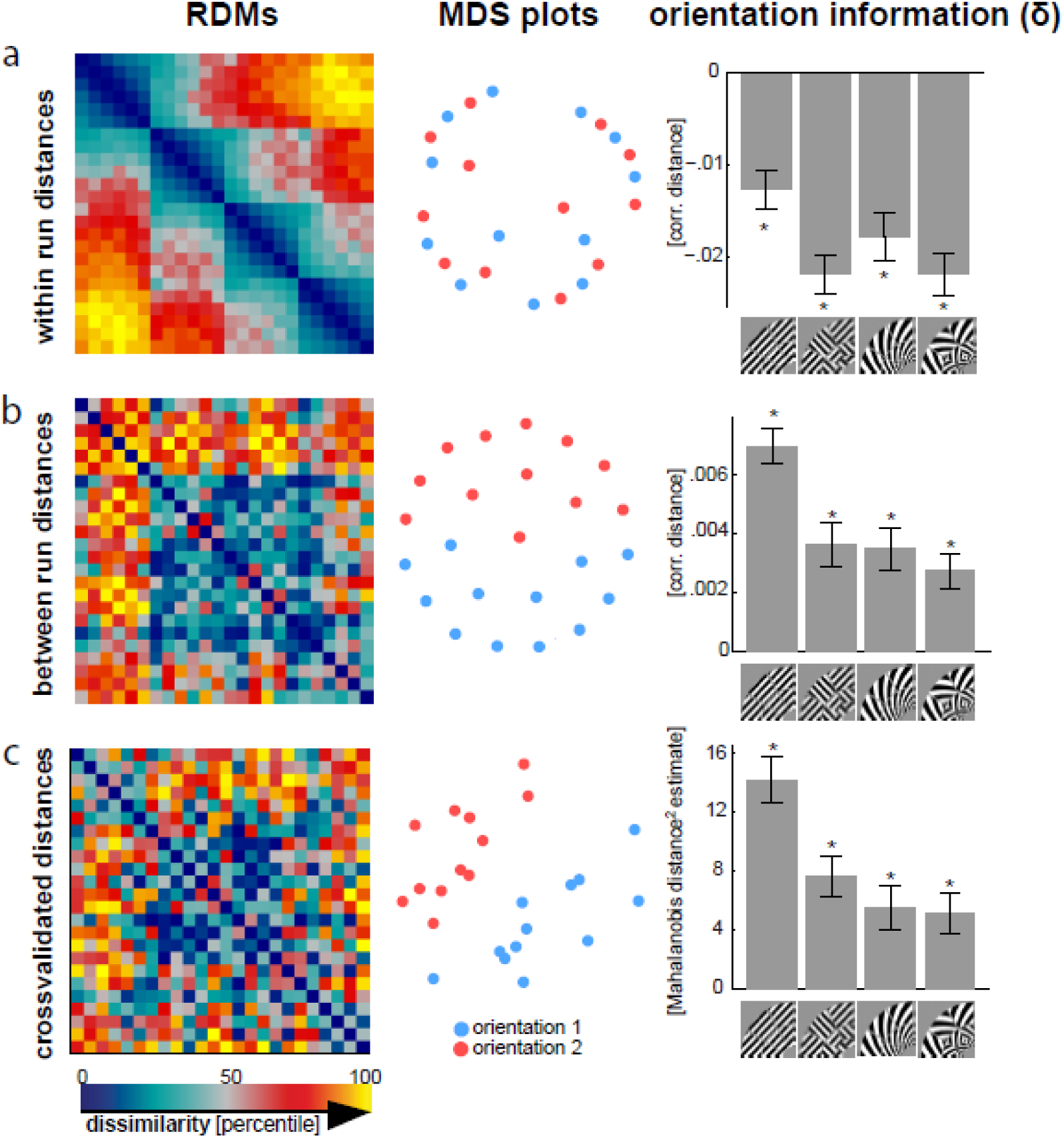
Crossvalidated distance estimates are unbiased by pattern drift. From left to right: a chronologically ordered RDM with all pairwise dissimilarities in a run, a multidimensional scaling plot and a bar graph depicting orientation information for each stimulus type. These are plotted based on: *a*. within-run pattern dissimilarities, b. between-run pattern dissimilarities and c. crossvalidated pattern dissimilarities.

### 3.6 Crossvalidation of fMRI pattern similarity estimates eliminates drift effects

One possible source of temporal pattern drift could be random walk like MRI signal fluctuations - physiological or scanner related - occurring within single voxel timecourses. Such fluctuations would lead to increased pattern dissimilarities for temporally distant patterns. This type of non-stationarity cannot be expected to be removed by voxelwise detrending or temporal highpass filtering because the trajectory of a random walk process is non-linear and has a high temporal frequency component. In addition, the direction of drifts evoked by a random walk process should be random. Therefore, if pattern drift is evoked by a random walk process then the drift effects should not replicate across independent observations (e.g. across runs). To test if this is the case, we determined whether fMRI pattern dissimilarities are robust to temporal drift if they are computed between independent data (see methods 2.5.2) or crossvalidated (see 2.5.3).

First, we constrained our analysis of pattern dissimilarities to between-run dissimilarities. This is similar to computing a crossvalidated distance estimate because the expected value of the estimates covariance between two fMRI patterns equals the true pattern covariance. This is because the correlation distance is computed between two independently measured fMRI patterns (i.e. coming from either run one or two). Therefore, error-components of fMRI patterns are expected to cancel out between them (for a complete explanation see section 2.5.2). Using between-run dissimilarities recovered fMRI pattern orientation information (average *δ* across subjects and stimuli was 0.0042; p<0.0001). Moreover relative orientation information across stimulus types (figure 4b-right) was found to be qualitatively similar to that obtained in our previous study using SVM classification (see Alink at al. 2013 and Fig 1A). This shows that computing dissimilarity estimates between independent fMRI runs is sufficient to restore orientation information.

Second, we computed crossvalidated squared Mahalanobis distance estimates (Walther et al, under revision) using leave-one-run-out crossvalidation for all within run pattern pairs (figure 4c, left). We chose Mahalanobis distance over correlation distance because crossvalidation may result in correlations outside the conventional [-1 1] boundaries (see section 2.5.3 and 7.1 in the appendix). Unlike the between-run distance, the crossvalidated Mahalanobis distance is an estimate of the true distance between orientation patterns and is truly ratio-scale with an interpretable zero point (Walther et al, under revision). Squared Mahalanobis distance estimates were found to be greater for between than within orientation pairs (average *δ* across subjects and stimuli was 8.12; p<0.0001) and relative orientation information across stimulus types (figure 4c-right) was found to be qualitatively similar to that obtained in our previous study using SVM classification (see Alink at al. 2013 and Fig 1A). This shows that crossvalidated distance estimates are unbiased by pattern drift.

## 4. Discussion

The main finding of this fMRI study is that response patterns are severely affected by temporal drift — pattern dissimilarity is shown to significantly increase as a function of temporal proximity of patterns. This effect occurs regardless of high-pass filtering and detrending of single voxel timecourses. For this particular dataset drift effects were confounded with stimulus orientation. As a consequence, orientation information could not be detected by comparing within-orientation pattern dissimilarities to between-orientation pattern dissimilarities. This exemplifies that temporal drift effects can obscure pattern effects of interest when pattern dissimilarity analysis is oblivious to drift-related pattern variance. Therefore, we propose that future studies analyzing fMRI pattern dissimilarities should account for such drift effects to increase the interpretability of results and the sensitivity to fMRI pattern effects of interest. We show here that this can be achieved both by drift modeling at the level of the representational dissimilarities and by means of crossvalidated distance measures.

Temporal drift was found to affect pattern dissimilarity in a consistent and predictable manner. Therefore, we were able to model the drift component in the pattern similarity structure and showed that the residual values contained significant orientation information. Our results suggest that this approach is effective in recovering drift-distorted effect. For our data a 5^th^ degree polynomial was required to model the drift precisely enough on the pattern dissimilarities. Whether this generalizes to other studies with different experimental designs and scanner parameters, however, remains to be shown.

Our data suggest that response-pattern dissimilarity estimates tend to increase with the temporal separation between the two stimuli. A possible cause for this effect could be random walk like fluctuations occurring within single voxel timecourses. These fluctuations cannot be expected to be removed by voxelwise detrending or temporal high-pass filtering because the trajectory of a random walk process is non-linear and has a high temporal frequency component. If a random walk process causes pattern drift than one should be able to eliminate drift effects by crossvalidating fMRI distance measures (Walther et al., under revision; also see sections 2.5.2 and A7.1) because the direction of a random walk process should not replicate across independent observations. Consistent with this prediction, our results indicate that crossvalidated fMRI distance estimates are drift-robust.

In sum, our results suggest that pattern drift effects can be successfully alleviated both by means of regressing these effects out and by using cross-validated distance estimates. We recommend using crossvalidating over the drift modeling approach because crossvalidation produces fully interpretable distance estimates that are unbiased by random noise in the fMRI patterns and have a meaningful zero point. Drift modelling, on the other hand, is an adhoc solution to drift effects and produces distance residuals that cannot be readily interpreted as dissimilarities anymore. However, the regressing out approach can be useful if one’s dataset does not allow for crossvalidation, e.g. if multiple imaging runs were not acquired or if conditions are not balanced across runs.

The fact that crossvalidation eliminates drift distortions in the RDMs suggest that random walk like fluctuations within single voxel time-courses might cause pattern drift. However, based on the current dataset we cannot tell whether these fluctuations represent time-continuous changes of brain states (Henriksson et al., 2015) or whether they can be attributed to scanner measurement artifacts. Future studies could clarify this issue by investigating the relationship between the temporal dynamics of pattern drift and fMRI scanning parameters. For example, one could test if pattern drift is accelerated when using fMRI sequences that cause greater heating of MRI gradient coils.

In this study differently oriented stimuli were presented in alternating fashion which led to a confound between stimulus orientation and drift effects. This confound could have been reduced by randomizing stimulus order. In general, we expect stimulus order randomization to significantly reduce the impact of drift effects on the outcome of pattern dissimilarity analyses and to remove drift effects as a systematic confound. However, given the magnitude of the drift effects, they may still significantly reduce the sensitivity of pattern dissimilarity analysis. Therefore, we recommend that future studies analyzing fMRI pattern dissimilarities both use a randomized stimulus sequence and account for drift effects during the analysis.

In summary, we have demonstrated that temporal drift has a prominent effect on fMRI patterns and that this effect can obscure pattern information about visual stimulus orientation. Pattern information, however, can be recovered by regressing out drift effects from pattern dissimilarities or by computing crossvalidated dissimilarity estimates. We recommend that future fMRI studies take pattern drift into account when analyzing pattern dissimilarities as this can greatly enhance the sensitivity to pattern effects of interest.

## 6. Acknowledgments

This work was supported by the UK Medical Research Council and by a European Research Council Starting Grant (261352) and Wellcome Trust Project Grant (WT091540MA) to NK, a Gates Cambridge Scholarship to AW and a British Academy postdoctoral fellowship to AA. JB is supported by The Ready Mind Project.

## 7. Appendix

### 7.1 Crossvalidated Pearson correlation estimate

Consider two mean-centered fMRI activity pattern of condition a of run one, û*_a_*, and condition b of run two, û*_b_*. Assume we are given two independent repetitions of each of a and b (e.g. from two functional fMRI runs), 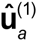, 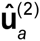 and 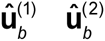, respectively. The fully crossvalidated Pearson correlation between a and b is then:

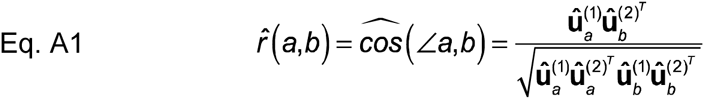

Note that unlike in the between-run correlation (Eq. 6), the variances of a and b are now computed using patterns from different runs. Again, we assume that each pattern estimate has a true underlying stimulus component and a noise pattern that is independent between runs (Eq. 4). Plugging the decomposed estimates into Eq. A1 then obtains as

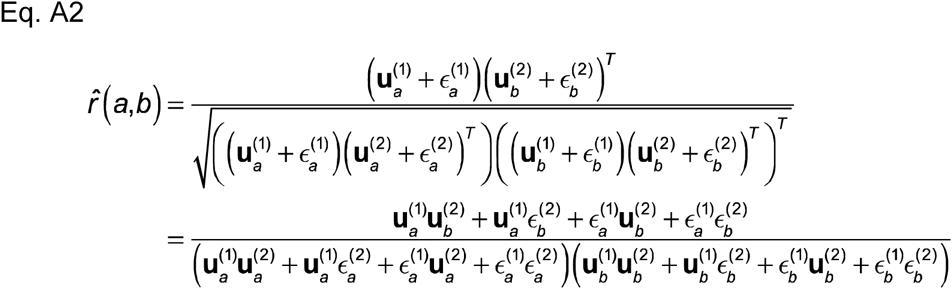

Since error terms from different runs are independent, the expected value of the fully crossvalidated *r*(*a,b*) is

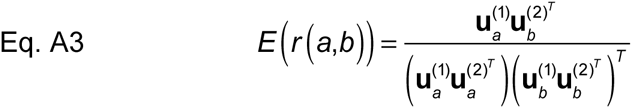

This value may exceed the [-1 1] range because the pattern variances in run 1 and 2 can be very different in scale and hence r may not meet the Cauchy-Schwarz inequality, i.e. 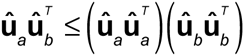.

## Supplementary materials

**Figure A1:**
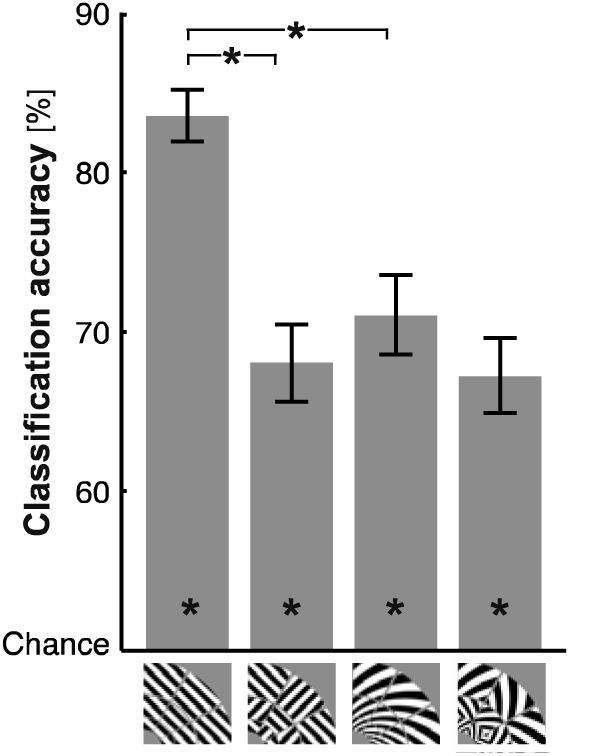
Stimulus identity is robustly decodable with linear support vector machine for all stimulus types. Average classification accuracies of the four stimulus types used in Alink et al. (2013): grating, spiral, and patch-swapped versions of both. Decoding was performed using a linear support vector machine (leave-one-subrun-out crossvalidation) for on multivariately noise-normalized V1 fMRI patterns. Error bars indicate standard error of the mean across 18 subjects. Asterisks on bars indicate above-chance classification accuracy (p < 0.01). Asterisks on horizontal brackets indicate significant difference (p < 0.01) between classification accuracies.

